# Inhibition of major histocompatibility complex-I antigen presentation by sarbecovirus ORF7a proteins

**DOI:** 10.1101/2022.05.25.493467

**Authors:** Fengwen Zhang, Trinity Zang, Eva M. Stevenson, Xiao Lei, Dennis C. Copertino, Talia M. Mota, Julie Boucau, Wilfredo F. Garcia-Beltran, R. Brad Jones, Paul D. Bieniasz

## Abstract

Viruses employ a variety of strategies to escape or counteract immune responses, including depletion of cell surface major histocompatibility complex class I (MHC-I), that would ordinarily present viral peptides to CD8+ cytotoxic T cells. As part of a screen to elucidate biological activities associated with individual SARS-CoV-2 viral proteins, we found that ORF7a reduced cell surface MHC-I levels by approximately 5-fold. Nevertheless, in cells infected with SARS-CoV-2, surface MHC-I levels were reduced even in the absence of ORF7a, suggesting additional mechanisms of MHC-I downregulation. ORF7a proteins from a sample of sarbecoviruses varied in their ability to induce MHC-I downregulation and, unlike SARS-CoV-2, the ORF7a protein from SARS-CoV lacked MHC-I downregulating activity. A single-amino acid at position 59 (T/F) that is variable among sarbecovirus ORF7a proteins governed the difference in MHC-I downregulating activity. SARS-CoV-2 ORF7a physically associated with the MHC-I heavy chain and inhibited the presentation of expressed antigen to CD8+ T-cells. Speficially, ORF7a prevented the assembly of the MHC-I peptide loading complex and causing retention of MHC-I in the endoplasmic reticulum. The differential ability of ORF7a proteins to function in this way might affect sarbecovirus dissemination and persistence in human populations, particularly those with infection- or vaccine-elicited immunity.

## Introduction

To replicate and propagate in a host population that presents an immunologically hostile environment, viruses typically employ a variety of strategies to escape or counteract immune responses. Severe acute respiratory syndrome coronavirus-2 (SARS-CoV-2), a member of the sarbecovirus subgenus, has been shown to antagonize the innate immune response through the action of viral proteins (1–3) and to escape humoral immunity through variation in the neutralizing epitopes of the spike protein (4–8). Evasion of cell-mediated immunity is accomplished by many viruses through the downregulation of surface expression of major histocompatibility complex-I, that would ordinarily present viral peptides to CD8+ cytotoxic T-cells (9–11). For example, HIV-1 renders the virus-infected cells less visible to CD8+ T-cells through Nef-induced endocytosis of MHC-I from the cell surface (12). In general, viruses from other families that are associated with chronic infections employ diverse mechanisms to deplete MHC-I from infected cell surfaces (10, 11). However, viruses associated with short-term, acute infection do not typically induce MHC-I downregulation.

The ~30-kb SARS-CoV-2 genome encodes structural proteins (E, M, N, and S), nonstructural proteins (nsp1 to nsp16), and several ‘accessory’ open reading frames (ORF3a, ORF6, ORF7a, ORF8, ORF10, ORF3b, ORF9b, and ORF9c) (13, 14). Analysis of coronavirus-host proteinprotein interactions, using affinity- or proximity-based approaches, has suggested that several viral proteins (ORF3a, ORF3b, ORF7a, ORF8, M, and nsp4) associate with host proteins that are enriched in endoplasm reticulum (ER) or Golgi, the organelles where viral peptides are loaded onto MHC-I molecule and transported to the cell surface for presentation to CD8+ T-cells (14, 15). Moreover, Some reports have indicated that SARS-CoV-2 ORF8 reduces expression of MHC-I on the surface of infected cells (16, 17). Here we show that SARS-CoV-2 ORF7a can inhibit antigen presentation by preventing the assembly of the MHC-I peptide loading complex and causing retention of MHC-I in the endoplasmic reticulum. Notably, ORF7a proteins from a sample of sarbecoviruses vary in their ability to induce MHC-I downregulation and a single-amino acid that is variable among sarbecovirus ORF7a proteins governs the differential ability to induce in MHC-I downregulation.

## Results

### SARS-CoV-2 ORF7a reduces cell surface MHC-I levels

To elucidate biological activities associated with individual SARS-CoV-2 viral proteins we used an HIV-1-based lentiviral vector (pSCRPSY) (18) to express each SARS-CoV-2 viral open reading frame, as annotated in (13, 14). Two days after transduction of human 293T cells, we measured MHC-I surface levels by flow cytometry using a pan-HLA class I-reactive monoclonal antibody. Expression of ORF7a reduced MHC-I levels on the cell surface by approximately 5-fold, whereas expression of other individual viral proteins (notably including ORF8 (16, 17)) had no effect on MHC-I surface levels (Figure 1). We also examined the impact of SARS-CoV-2 ORFs on the expression of tetherin, a cell surface antiviral protein that traps enveloped virions from various virus families that bud through cell membranes. None of SARS-CoV-2 viral ORFs reduced the levels of tetherin stably expressed on the surface of 293T cells (Figure S1), underscoring the specificity of the effect of ORF7a on MHC-I. Of note, expression of two viral proteins (nsp1 and ORF6) was not accomplished in our screen, as lentiviral vectors encoding these proteins were low titer, in line with the previous findings that nsp1 suppresses host protein synthesis (19) and that ORF6 interferes with nuclear transport machinery (14, 15).

**Figure 1.**
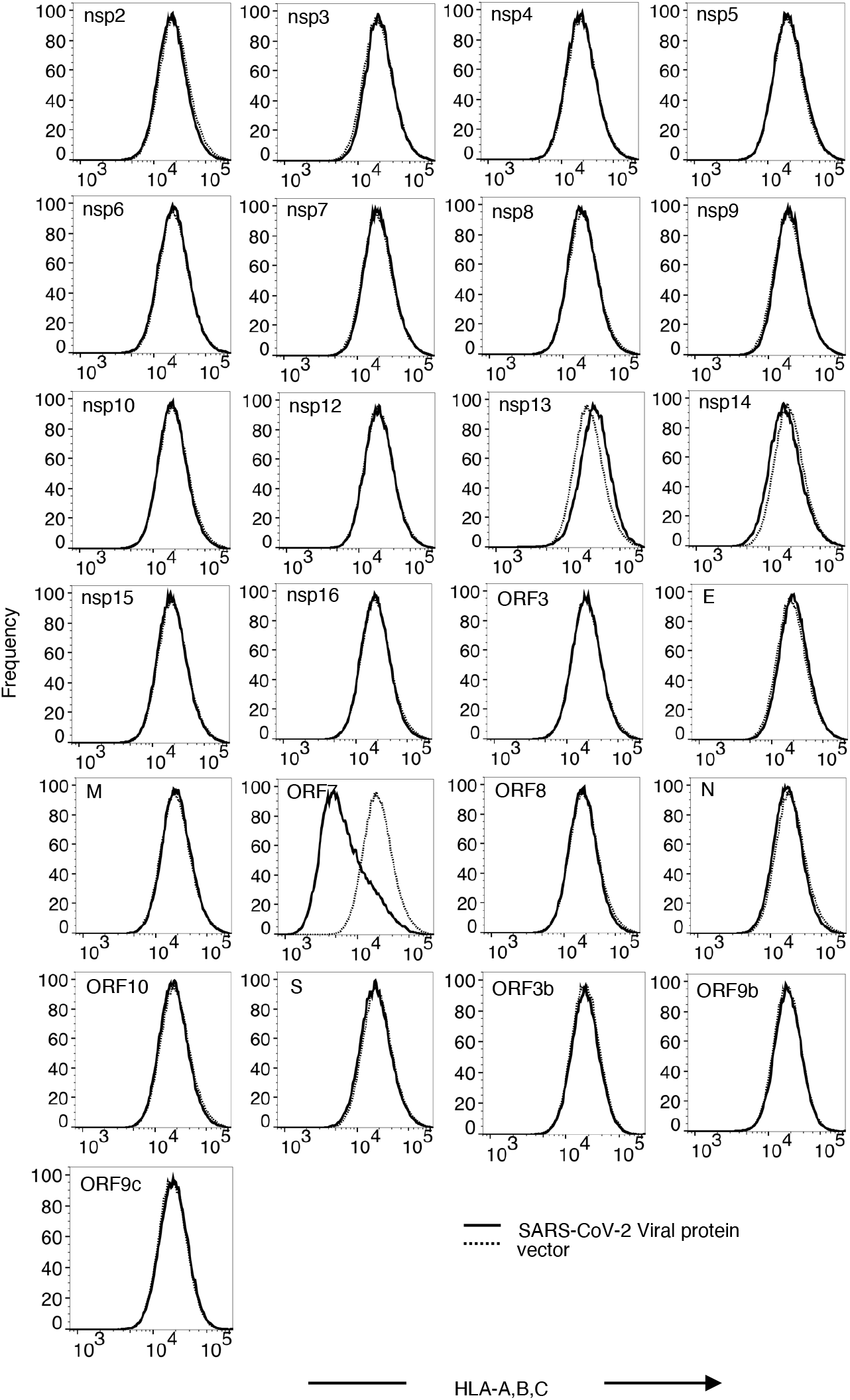
Effect of SARS-COV-2 ORFs on MHC-I cell surface levels in 293T cells. Human 293T cells transduced with SCRPSY-based lentiviral vector expressing individual SARS-CoV-2 viral proteins, at an MOI of 0.5 and cell surface MHC-I detected with a pan-HLA-ABC antibody (W6/32) and flow cytometry two days later. Cells transduced (TagRFP+ population) with viral protein expression vector (solid line) and empty lentiviral vector (dotted line) were gated and compared. Representative of three experiments.

ORF7a caused reduced MHC-I cell surface levels in other human cells such as Huh7.5 and U2OS (Figure 2A) suggesting that its activity is not cell type-specific. Because the recognition of MHC-I by the W6/32 antibody could be influenced by association between heavy chains (HC) and β2-microglobulin (β2M) (20, 21), we next confirmed that downregulation occurred, as assessed with a different antibody, specific for the HLA-A HC (Figure 2B). MHC-I surface levels were also depleted following SARS-CoV-2_USA-WA1/2020_ infection of A549/ACE2 cells, specifically in the infected nucleocapsid-positive subpopulation (Figure 2C, D). However, MHC-I downregulation was largely maintained in cells infected with SARS-CoV-2 lacking ORF7a, suggesting the existence of additional, redundant means of MHC-I downregulation (Figure 2C, D). We did not observe the previously reported MHC-I downregulation induced by ORF8 expression, nor expression of other individual SARS-CoV-2 ORFs (Figure 1), suggesting that additional mechanisms of MHC-I downregulation requires the concerted action of multiple SARS-CoV-2 proteins.

**Figure 2.**
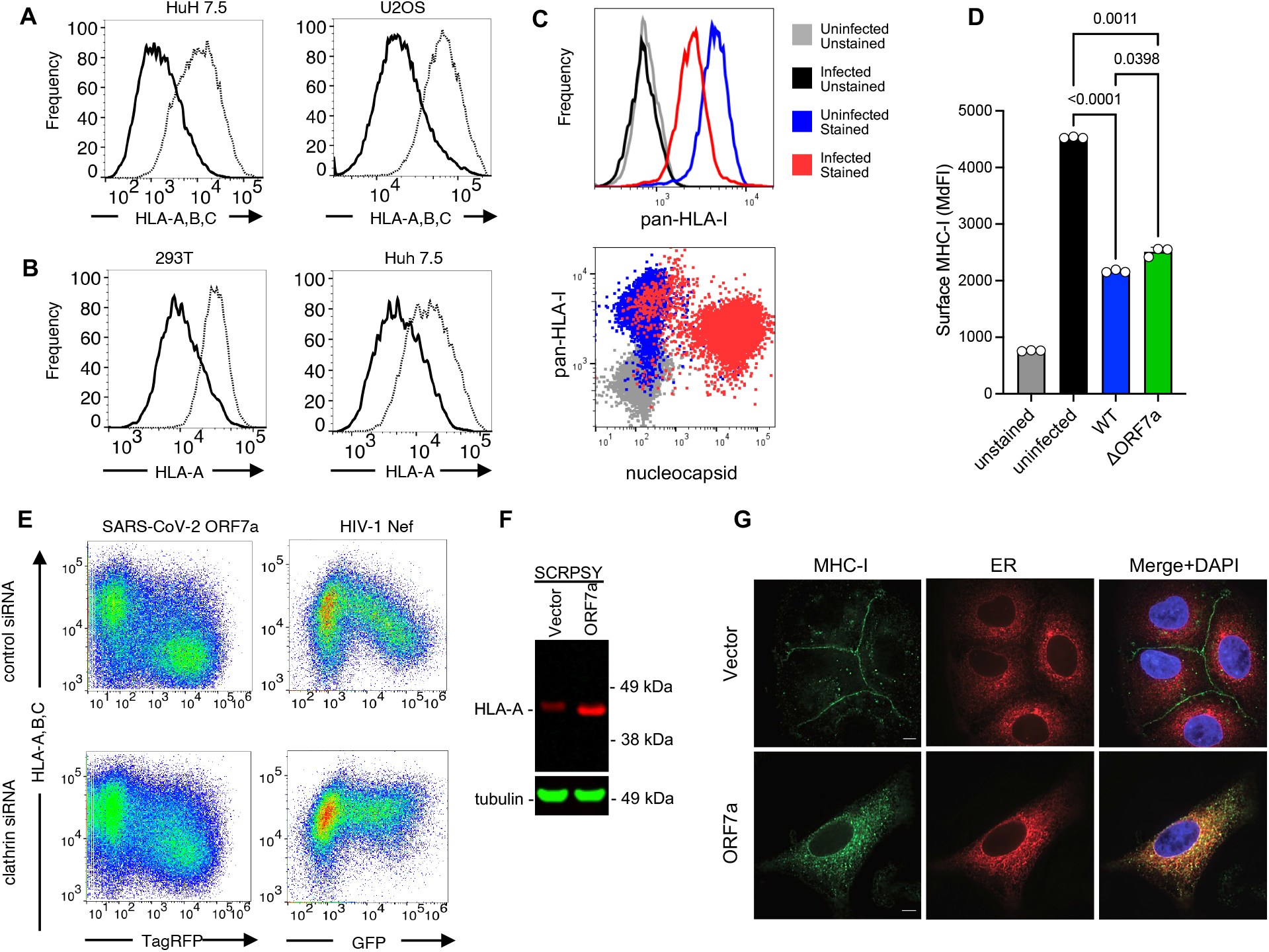
MHC-1 downregulation by ORF7a. (A) HuH7.5 cells, U2OS cells or 293T cells transduced with pSCRPSY-based lentiviral vector expressing SARS-CoV-2 ORF7a at an MOI of 0.5. and stained with a pan-HLA-ABC antibody (W6/32) or an HLA A-allele specific antibody (YTH 862.2) followed by flow cytometry. Cells transduced (TagRFP+ population) with the ORF7a expression vector (solid line) and empty lentiviral vector (dotted line) were gated and compared. Representative of three experiments. (C) A549/ACE2 cells stained with stained with a pan-HLA-ABC antibody (W6/32) and SARS-CoV-2 nucleocapsid antibody after infection with SARS-CoV-2_USA-WA1/2020_ (D) MHC-I surface levels (Median fluorescent intensity (MdFI)) in A549/ACE2 cells stained with a pan-HLA-ABC antibody (W6/32) after infection with SARS-CoV-2_USA-WA1/2020_ (WT) or icSARS-CoV-2-mNG (ΔORF7). (E) Flow cytometric analysis of 293T cells with a pan-HLA-ABC antibody (W6/32) following transduction with a pSCRPSY-lentiviral vector expressing SARS-CoV-2 ORF7a (left panels) or a lentivirus vector expressing Nef and GFP (right panels), at an MOI of 0.5. Cells were transfected with control (upper panels) or clathrin HC-depleting (lower panels) siRNA prior to transduction. Representative of three experiments. (F) Western blot analysis of 293T cells 48h after following transduction with empty (vector) or SARS-CoV-2 ORF7a expressing SCRPSY with anti-HLA-A(red) and anti Tubulin (green). (G) Immunofluorescent staining of A549 cells with anti HLA-A (green) following transduction with empty SCRPSY, (vector) or ORF7a -expressing SCRPSY (ORF-7a) and ER-GFP BacMam (red) a single optical section following deconvolution microscopy is shown.

After loading with peptides in the endoplasmic reticulum (ER), MHC-I molecules are transported via the Golgi to the plasma membrane and are subsequently turned over by endocytosis (22, 23). To begin to elucidate how ORF7a might reduce cell surface MHC-I levels, we first depleted clathrin, with small interfering RNA and found that clathrin knockdown, as reported previously (24), inhibited the downregulation of MHC-I by Nef (Figure 2E). Conversely, clathrin depletion had only a minor effect on MHC-I downregulation by ORF7a, suggesting that ORF7a disturbs surface MHC-I levels in a distinct, clathrin-independent manner (Figure 2E). Western blot analysis revealed no deficit in the total amount of MHC-I in cells expressing ORF7a. Indeed, when cells were transduced with the ORF7a lentivirus at sufficient MOI that most cells expressed ORF7a, the total amount of endogenous cellular MHC-I was increased, and displayed slightly accelerated migration, suggesting that ORF7a induces intracellular accumulation and altered post-translational modification of MHC-I (Figure 2F).

Immunofluorescent staining of endogenous MHC-I in A549 cells showed that ORF7a profoundly altered the subcellular distribution of MHC-I molecules (Figure 2G). Specifically, ORF7a induced a reduction in of MHC-I HC fluorescence that could be observed at the cell surface, consistent with the flow cytometric analysis (Figure 2A), while intracellular MHC-I HC accumulated (Figure 2G), The accumulated intracellular MHC-I HC was partly colocalized with a marker of the ER (Figure 2G). The MHC-I component β2M was similarly redistributed to intracellular locations upon ORF7a expression, and partly colocalized with ORF7a (Figure S2). We conclude that SARS-CoV-2 ORF7a blocks the ability of MHC-I to move through the secretory pathway to the cell surface.

### Determinants of sarbecovirus ORF7a MHC-I downregulation activity

We tested whether ORF7a proteins from a panel of bat SARS-related coronaviruses (SARSr-CoV), namely BM48-31, HKU3-1, Rf1, Rs672, ZC45, ZXC21, and RaTG13 shared the ability to reduce cell surface MHC-I levels (25). The RaTG13, Rf1, ZC45, and ZXC21 ORF7a proteins, but not those from BM48-31, HKU3-1, and Rs672 ORF7, reduced MHC-I surface levels (Figure 3A). Western blot and immunofluorescence analyses showed that the sarbecovirus ORF7a proteins were expressed in transduced cells at a similar level, and those that induced surface downregulation also induced intracellular accumulation and altered migration of MHC-I (Figure 3B, Figure S3).

**Figure 3.**
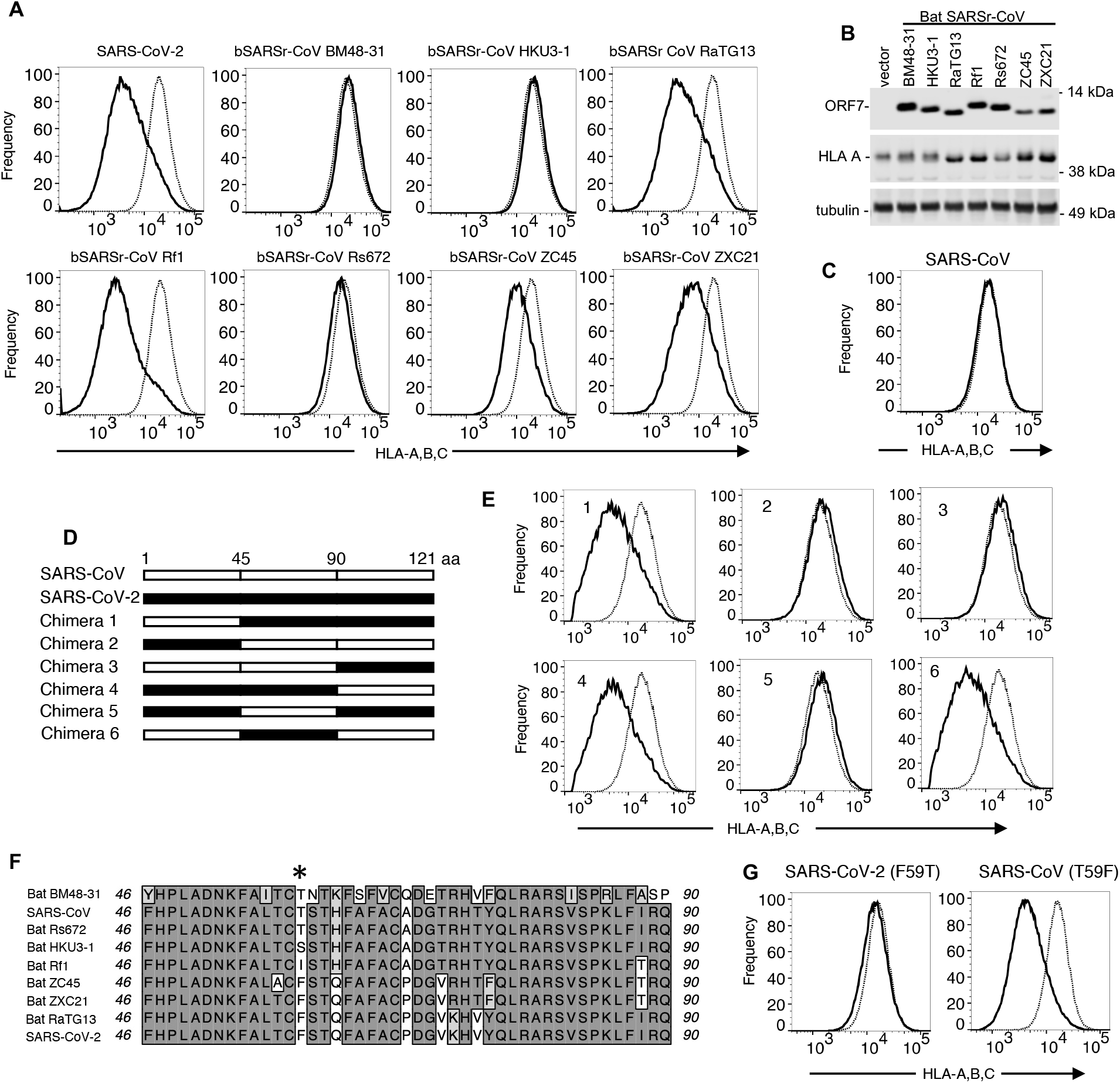
Determinants of MHC-I downregulation by sarbecovirus ORF7a proteins. (A) Flow cytometric analysis of 293T cells with a pan-HLA-ABC antibody (W6/32) following transduction with a pSCRPSY-lentiviral vector expressing ORF7a proteins from SARS-CoV-2 bat SARSr coronaviruses BM48-31/BGR/2008 (YP_003858589.1), HKU3-1 (AAY88871.1), RaTG13 (QHR63305.1), Rf1/2004 (ABD75319.1), Rs_672/2006 (ACU31037.1), ZC45 (AVP78036.1) and ZXC21 (AVP78047.1). Cells transduced (TagRFP+ population) with the ORF7a expression vector (solid line) and empty lentiviral vector (dotted line) were gated and compared. Representative of three experiments. (B) Western blot analysis of 293T cell lysates following transduction with SCRPSY with no ORF7a (vector) or the ORF7a proteins from bat SARS-like coronaviruses as in (A) (C) Flow cytometric analysis of 293T cells with a pan-HLA-ABC antibody (W6/32) following transduction with a pSCRPSY-lentiviral vector expressing SARS-CoV. Cells transduced (TagRFP+ population) with the ORF7a expression vector (solid line) and empty lentiviral vector (dotted line) were gated and compared. (D) Schematic representation of chimeras generated between SARS-CoV and SARS-CoV-2 ORF7a proteins. (E) Flow cytometric analysis of 293T cells with a pan-HLA-ABC antibody (W6/32) following transduction with a pSCRPSY-lentiviral vector expressing chimeric ORF7a proteins shown in (D) Cells transduced (TagRFP+ population) with the ORF7a expression vector (solid line) and empty lentiviral vector (dotted line) were gated and compared. (F) Amino acid alignment of SARS-CoV, SARS-CoV-2, bat SARSr-CoV ORF7a proteins. Identical residues are shaded. Star indicates the amino acid at position 59 that were mutated in proteins used in (D). (G) Flow cytometric analysis of 293T cells with a pan-HLA-ABC antibody (W6/32) following transduction with a pSCRPSY-lentiviral vector expressing point mutant ORF7a proteins. Cells transduced (TagRFP+ population) with the ORF7a expression vector (solid line) and empty lentiviral vector (dotted line) were gated and compared.

Notably, the ORF7a protein from SARS-CoV was among those that did not affect MHC-I surface levels (Figure 3C), despite encoding 85% identical amino acids to SARS-CoV-2 ORF7a. Moreover, unlike SARS-CoV-2 ORF7a, and the active bat SARSr-CoV ORF7a proteins, the SARS-CoV ORF7a protein showed little tendency to colocalize with MHC-I (Figure S4). To map the determinants of the differential ability to modulate cell surface MHC-I levels, we constructed 6 chimeric SARS-CoV-2/SARS-CoV ORF7a expression vectors (Figure 3D). Chimeric ORF7a proteins containing the central region (residues 46-90) from SARS-CoV-2 all downregulated MHC-I from the cell surface while those encoding the equivalent region from SARS-CoV did not (Figure 3E). Among the sarbecovirus ORF7a proteins, the presence of hydrophobic residues (F or I) at position 59 within this region correlated with the ability to downregulate MHC-I, while the presence of a polar residue (T or S) correlated with inactivity (Figure 3F). Thus, to test the importance of the amino acid at position 59, we introduced an F59T substitution into SARS-CoV-2 ORF7a and found this substitution abolished MHC-I downregulating activity, while the reciprocal substitution in SARS-CoV ORF7a (T59F) substitution led to acquisition of MHC-I downregulating activity (Figure 3G). We conclude that residue 59, a variable position among sarbecovirus ORF7a proteins, is a critical determinant of their ability to modulate cell surface MHC-I levels.

### Interaction between MHC-I and ORF7a

To test whether ORF7a proteins could physically associate with MHC-I, we immunoprecipitated the endogenously expressed MHC-I HC, in ORF7a expressing cells. The SARS-CoV-2 ORF7a protein could be co-immunoprecipitated from transduced cells with an anti-HLA-A antibody, and the amount of co-immunoprecipitated ORF7a but not the total level of ORF7a protein was reduced to background levels by the functionally inactivating F59T substitution in SARS-CoV-2 ORF7a (Figure 4A). Conversely, SARS-CoV ORF7 was poorly co-immunoprecipitated by the anti-HLA-A antibody but the T59F gain of function substitution increased the amount of co-immunoprecipitated ORF7a (Figure 4A). In reciprocal immunoprecipitation experiments, HLA-A could be immunoprecipitated by an ORF7a antibody from cells expressing wild-type SARS-CoV-2 ORF7a or SARS-CoV ORF7a(T59F) mutant (Figure 4B). Conversely, expression of the SARS-CoV-2 ORF7a(F59T) mutant or wild-type SARS-CoV ORF7a resulted in less efficient coprecipitation of the HLA-A protein (Figure 4B). We conclude that ORF7a associates with MHC-I HC and that single amino-acid substitutions in ORF7a that confer or ablate MHC-I downregulating activity simultaneously confer or ablate the ability of ORF7a to coimmunoprecipitate with MHC-I.

**Figure 4.**
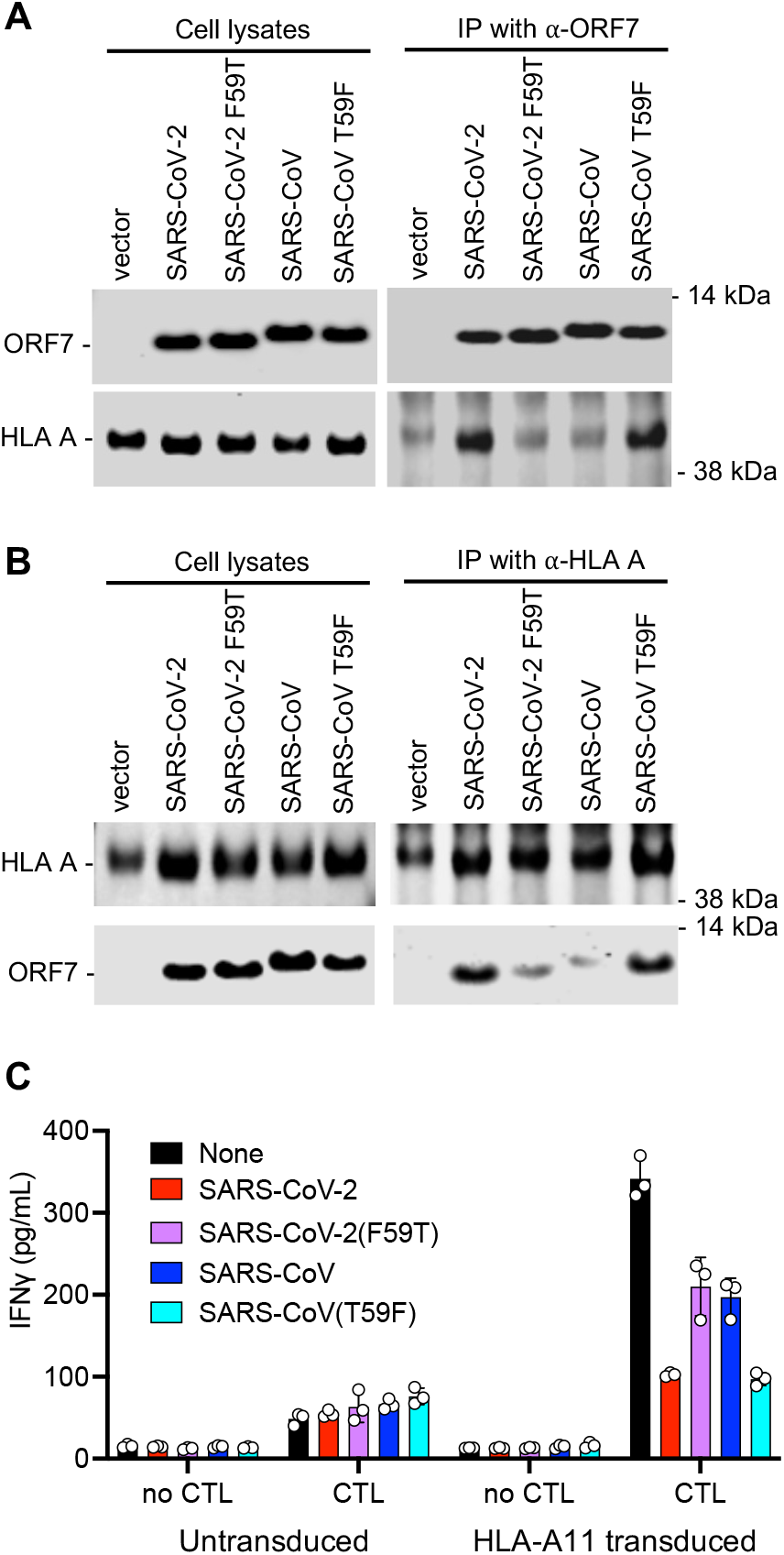
ORF7a physically associated with MHC-I and inhibits antigen presentation. (A, B) Western blot analysis of cell lysates and immunoprecipitates (IP) from 293T cells transduced with SCRPSY expressing no ORF7a (vector) or the indicated wild-type (WT) or mutant ORF7a proteins. Immunoprecipitation was carried out with mouse anti-SARS-CoV ORF7a mAb (3C9) precipitated ORF7a and MHC-I proteins were detected with rabbit anti-ORF7a and an HLA A-specific antibody (EP1395Y). (A) Alternatively, immunoprecipitation was carried out with a mouse HLA class I ABC monoclonal antibody (66013-1-Ig) and precipitated ORF7a and MHC-I proteins were detected with rabbit anti-ORF7a antibody and an HLA A-specific antibody (EP1395Y), respectively. Representative of three experiments. (C) Levels of IFN-γ in supernatants following 16h of incubation of the AK11 CD8+ T-cell clone with 293T target cells that were either untransduced or transduced with an HLA-A11 expression vector, followed by an ORF7a expressing (or control) SCRPSY vector, and then transfected with an HIV-1 Gag expression plasmid.

### ORF7a inhibits MHC-I antigen presentation to a CD8+ T-cell clone

To determine whether SARS-CoV-2 ORF7a could inhibit antigen presentation, we engineered 293T cells to express HLA-A11 and then expressed an HIV-1 Gag protein and an ORF7a protein therein. We used these HIV-1 Gag-expressing cells as ‘targets’ for an HLA-A11 restricted CD8+ T-cell clone that recognizes the HIV-1 Gag peptide ACQGVGGPSHK (AK11) and measured the ability of the CD8+ T-cell clone to respond to the Gag-expressing target cells by secreting IFNγ. The AK11 CD8+ T-cell clone indeed responded to Gag-expressing 293T cells by secreting IFNγ, and this response was inhibited, almost to background levels, when the 293T target cells also expressed SARS-CoV-2 ORF7a (Figure 4C). The loss of function (F59T) SARS-CoV-2 ORF7a mutant and the wild-type SARS-CoV ORF7a proteins were less potent in reducing AK11 CD8+ T-cell clone responsiveness. Conversely, the SARS-CoV gain of function (T59F) mutant reduced the IFNγ response as potently as the WT SARS-CoV-2 ORF7a protein (Figure 4C).

### Disruption of the MHC-I peptide loading complex and ER retention induced by ORF7a

We next attempted to determine more precisely how ORF7a interferes with MHC-I transport through the secretory pathway, and with antigen presentation at the cell surface. As MHC-I moves through the Golgi, a single N-linked glycan on the HC is remodeled from an initially endoglycosidase H (Endo H) sensitive-high mannose form to a complex Endo H resistant form(26). Western blot analysis of Endo H treated cell lysates from control A549 cells indicated that at steady state the majority of MHC-I HC glycan was Endo H resistant, while in a cell population in which the cells expressed ORF7a, about 50% the HLA-A HC was Endo H sensitive (Figure 5A). The inactive SARS-CoV-2(F59T) and WT SARS-CoV proteins did not induce the appearance of the Endo H-sensitive species while the SARS-CoV(T59F) gain of function mutant had acquired this activity. Moreover, the abilities of members of the panel of bat SARSr-CoV proteins to deplete MHC-I proteins from the cell surface correlated with their abilities to induce the accumulation of Endo H-sensitive MHC-I HC species (Figure 5A). We conclude that the sarbecovirus ORF7a which cause depletion of cell surface MHC-I do so by inducing retention of MHC-I in pre-medial Golgi compartment(s).

**Figure 5.**
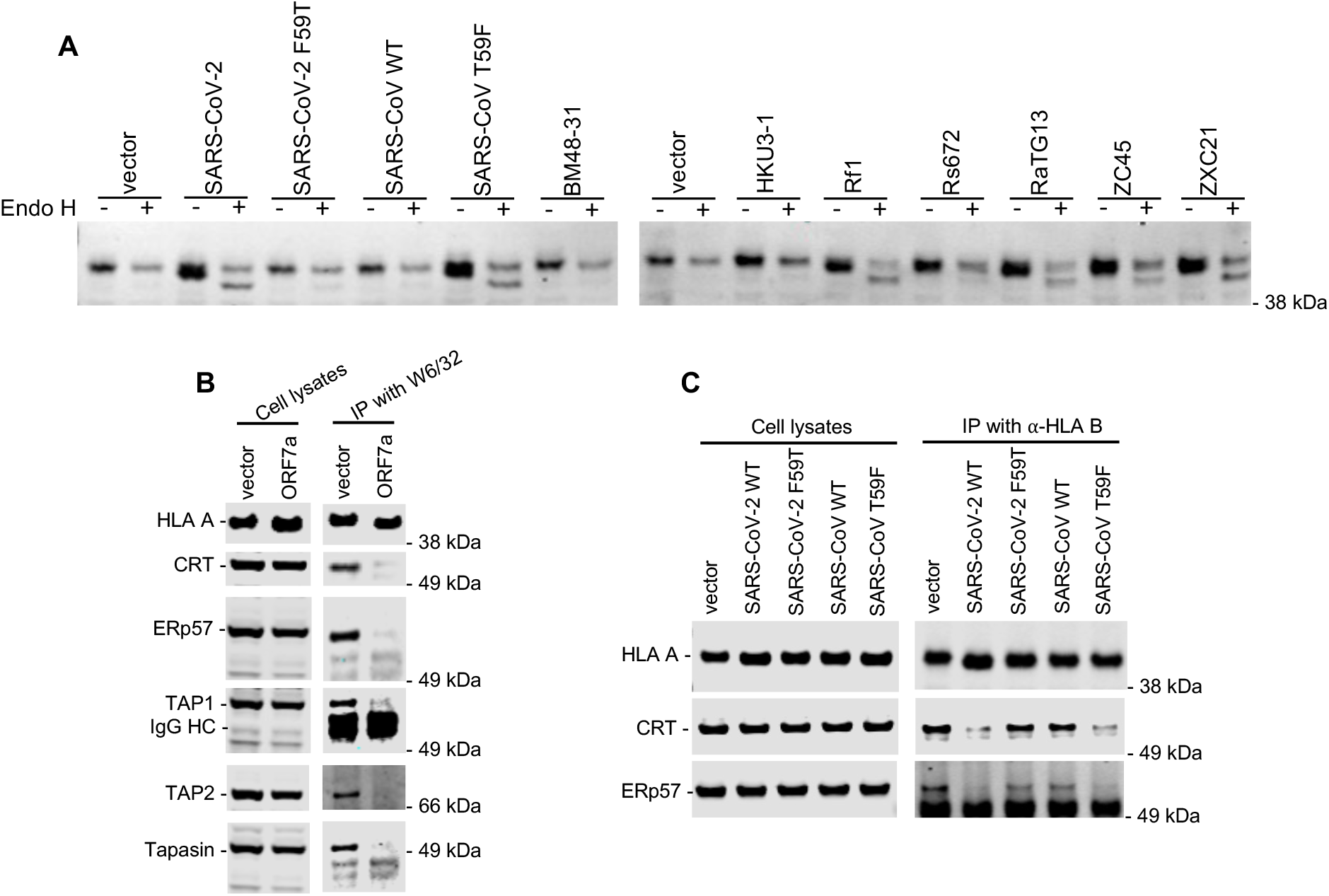
ER retention and disruption of the MHC-1 peptide loading complex assembly by ORF7a. (A) Western blot analysis of cell lysates of A549 cells transduced with SCRPSY expressing no ORF7a (vector) or the ORF7a proteins from SARS-CoV-2, SARS-CoV (or mutants thereof) or ORF7a proteins from the indicated bat SARSr-CoVs at an MOI of 0.5. Cell lysates were untreated or treated with endoglycosidase H, as indicated prior to analysis. (B) Western blot analysis of cell lysates and immunoprecipitates (IP) from 293T cells transduced with SCRPSY expressing no ORF7a (vector) or the SARS-CoV-2 ORF7a protein. Immunoprecipitation was carried out with a pan-HLA-ABC antibody (W6/32). Immunoprecipitated proteins were detected with rabbit anti-Calreticulin antibody (CRT), anti-ERp57 antibody, anti-TAP1 antibody, anti-TAP2 antibody and anti-Tapasin antibody. Representative of three experiments. (C) Western blot analysis of cell lysates and immunoprecipitates (IP) from 293T cells transduced with SCRPSY expressing no ORF7a (vector) or the indicated WT and mutant ORF7a proteins. Immunoprecipitation carried out with a rabbit HLA-B specific antibody (PA5-35345). Immunopreciptated proteins were detected with mouse anti-Calreticulin antibody and anti-ERp57 antibody respectively. Representative of three experiments.

For antigen presentation, peptides are loaded into MHC-I in the ER through an interaction of MHC-I with the peptide loading complex (PLC). The PLC includes the transporter associated with antigen processing (TAP) proteins, TAP1 and TAP2, as well as chaperones/cofactors Tapasin, ERp57 and calreticulin that each bind directly or indirectly to the MHC-HC/ β2 microglobulin complex in the ER lumen(9). Western blot analysis of proteins that co-immunoprecipitated with endogenous MHC-I revealed that expression of SARS-CoV-2 ORF7a blocked the co-immunoprecipitation of each the aforementioned MHC-I associated proteins with MHC-I HC without affecting the steady state levels of these proteins. To confirm the specificity of these effects, we employed the SARS-CoV-2 loss of function (F59T) and SARS-CoV gain of function (T59F) mutants. In similar co-immunoprecipitation assays in which endogenous HLA-B was immunoprecipitated, the co-immunoprecipitation of calreticulin and ERp57 was inhibited by the active WT SARS-CoV-2 and SARS-CoV(T59F) ORF7a proteins, but not by the inactive SARS-CoV-2(F59T) and WT SARS-CoV ORF7a proteins. We conclude that active ORF7a proteins disrupt the assembly of the MHC-I PLC in the ER and prevents the movement of peptide loaded MHC-I complexes to the cell surface for antigen presentation.

## Discussion

Inhibition of antigen presentation through the downregulation of MHC-I peptide complexes is a long-recognized means by which certain viruses mitigate the inhibitory action of immune responses on their replication (10). MHC-I downregulation is frequently associated with viruses that cause persistent or chronic infection, or viruses with large DNA genomes and many immunomodulatory genes (11). Such downregulation is less frequently associated with RNA viruses or those that cause acute, rapidly resolving infections (11). Nevertheless, by reducing the density of viral peptides that are displayed on the surface of infected cells, viruses that cause acute infections such as SARS-CoV-2 could gain a competitive advantage through MHC-I downregulation. This advantage would be most pronounced in contexts where pre-existing T cell responses could limit virus replication, such as re-infection, and might also extend to limiting the protective effects of pre-existing T cell responses to epitopes that are conserved in distinct coronaviruses (27). Moreover, while SARS-CoV-2 is typically viewed as an acute respiratory infection, it is increasingly recognized that SARS-CoV-2 infection can persist for months in certain anatomical sites (e.g. the gut) or in individuals with sub-optimal immune responses (28–30). Moreover, the natural history of sarbecovirus infection in non-human species such as bats, and the degree to which it is a persistent infection is poorly documented. Thus, there are several reasons to think that MHC-I downregulation might be an evolved property in sarbecoviruses. Some recent studies have suggested that SARS-CoV-2 ORF8 protein induces MHC-I downregulation(16, 17). We did not observe such an effect. Nevertheless, an ORF7a-deleted SARS-CoV-2 induced MHC-I downregulation almost as efficiently as did an ORF7a-intact virus. This finding suggests that additional redundant mechanisms are employed by SARS-CoV-2 to deplete surface MHC-I and reduce visibility to responding CD8+ T-cells. For example, a recent study has suggested that SARs-CoV-2 ORF6 inhibits transcriptional upregulation of MHC-I in response to infection (31).

The variation among the ability of bat SARSr-CoV ORF7a proteins to reduce surface MHC-I levels is intriguing. It is unclear whether MHC-I downregulation is a ‘natural’ function of ORF7a in bat hosts or whether its ability to downregulate MHC-I in humans is fortuitous. The structure of the ORF7a protein resembles an immunoglobulin domain, and position 59 is a surface exposed position. Variation at this critical determinant of MHC-I downregulation could reflect genetic conflict with variable MHC-I proteins in animal hosts, and lack of historical adaptation to humans. In any case, the differential ability of members of this pandemic threat virus subgenus to manipulate human MHC-I molecules could impact their ability to evade CD8+ T cell immunity elicited by prior sarbecovirus infection or vaccination, with substantial potential to affect human health.

## Acknowledgments

This work was supported by a grant from the National Institutes of Allergy and Infectious Diseases P01 AI165075 (to PDB) UM1 AI164565, R01 AI147845 (to RBJ). and by the Howard Hughes Medical Institute. The Ragon BSL3 laboratory is supported by the Harvard CFAR (P30 AI060354).

## Author contributions

FZ, JB, WGB, RBJ, PDB conceived and designed the experiments. TZ did the immunofluorescence experiments, XL assisted with plasmid construction, JB and WGB did the live SARS-CoV-2 experiments. EMS, DCC, and TMM did the antigen presentation experiments. FZ did all the other experiments. FZ and PDB wrote the paper with input for all the other authors.

## Materials and Methods

### Antibodies

Primary antibodies, including anti-ORF7a, anti-pan HLA ABC, HLA allele-specific antibodies, and peptide-loading complex antibodies, are listed and described in the Supplementary Information Table. Secondary antibodies included goat anti-rabbit and goat anti-mouse from Novus conjugated to DyLight 650 or Janelia Fluor 646 and goat anti-mouse from ThermoFisher Scientific conjugated to Alexa Flour 488 for immunofluorescence, or IRDye® 800CW, or IRDye® 680 (LI-COR Biosciences) for Western blot analysis.

### Plasmid Construction

All SARS-CoV-2 viral proteins, including non-structural proteins (NSPs), open-reading frames (ORFs) and structural proteins (E, M, N and S) annotated in the viral genome (GenBank accession MN985325 (13)) and (14)) were human codon-optimized using GenSmart™ Codon Optimization and synthesized by IDT DNA Technologies as gBlocks. ORF7a genes from SARS-CoV and the various bat SARSr-CoVs were synthesized based on their amino acid sequences, which included SARS coronavirus Tor2 (GenBank AAP41043.1), bat coronavirus BM48-31/BGR/2008 (YP_003858589.1), bat SARS coronavirus HKU3-1 (AAY88871.1), bat SARS CoV Rf1/2004 (ABD75319.1), bat SARS coronavirus Rs_672/2006 (ACU31037.1), bat coronavirus RaTG13 (QHR63305.1), bat SARS-like coronavirus ZC45 (AVP78036.1) and bat SARS-like coronavirus ZXC21 (AVP78047.1). The synthesized nucleotide sequences are listed in the Supplementary Table 2. For each gene fragment, XhoI and NotI sites were added at 5’ end and 3’ end, respectively and a typical kozak sequence (GCCACC) was inserted before each start codon (ATG). The synthetic genes were digested with XhoI and NotI and inserted into an HIV-1-based lentiviral expression vector (pSCRPSY) which also expresses a puromycin-resistance cassette and TagRFP.

To make the SARS-CoV (T59F) or SARS-CoV-2 ORF7a (F59T) expression vectors, overlap-extension PCR amplification were performed with primers that incorporated the corresponding nucleotide substitutions using corresponding wild-type plasmids as template, respectively. After digestion with XhoI and NotI, the purified PCR products were then inserted into the pSCRPSY expression vector. HLA A11 gene fragment was PCR amplified and, after digestion with XhoI and NotI, subcloned into a retroviral vector (LMNi-bsd).

### Cell lines

The human embryonic kidney HEK-293T cell line, human hepatoma-derived HuH-7.5 cell line, and human alveolar basal epithelial A549 cell line were maintained in DMEM supplemented with 10% fetal bovine serum (Sigma F8067) and gentamicin (Gibco). Human bone osteosarcoma epithelial cells (U2OS, ATCC® HTB-96™) were grown in McCoy’s 5a Medium Modified (ATCC® 30-2007™) /10%FCS/gentamicin. All cell lines used in this study were monitored by SYBR® Green real-time PCR RT assay periodically to ensure the absence of retroviral contamination and were stained with DAPI to test for mycoplasma contamination.

### Assessment of MHC-I downregulation by ORF7a

To assess HLA downregulation by SARS-CoV viral proteins, viral stocks were generated by transfecting 5 μg of pSCRPSY-based expression plasmids encoding either no SARS-CoV viral protein (empty vector) or individual SARS-CoV open reading frame, 5 μg of an HIV-1 Gag-Pol expression plasmid (pCRV1/GagPol) and 1 μg of VSV-G expression plasmid in 293T cells in 10-cm dishes using polyethylenimine (PolySciences). Virus-containing supernatant was collected and filtered (0.2μm) 2 days later. The viral stocks were then used to inoculate10^5^ target cells (293T, U2OS, or HuH7.5) in 12-well plates at an MOI of 0.5. At 48 h post-transduction, cells were detached from plates with 5 mM EDTA in PBS and stained for cell surface HLA-A, B, C expression with anti-HLA-A, B, C antibody conjugated to AF647 (W6/32, Biolegend). In some experiments, cell surface HLA-A level was determined with antibody against the HLA-A allele (YTH862.2), followed by Goat anti-Rat Alexa Fluro 488. The same procedure was done to quantify tetherin downregulation, except that 293T cells stably expressing HA-tagged tetherin were used as target cells, and cells were stained with anti-HA antibody. Flow cytometric analysis was performed using Attune® NxT Acoustic Focusing Cytometer (ThermoFisher Scientific).

To assess the effect of clathrin knockdown on MHC-I downregulation by SARS-CoV-2 ORF7a or HIV Nef, 293T cells were seeded one day prior to transfection with clathrin siRNA (Horizon Discovery, Dharmacon™) using Invitrogen Lipofectamine RNAiMax (ThermoFisher Scientific). One day later, the cells were transduced with lentivirus virus stocks expressing SARS-CoV-2 ORF7a or HIV Nef and harvested for FACS analysis 48 hours later.

### Assessment of MHC-I downregulation after live SARS-CoV-2 infection

Viral stocks of USA-WA1/2020 (BEI Resources) and icSARS-CoV-2-mNG (UTMB WRCEVA (32)) were generated by expanding virus on Vero-E6 cells (BEI Resources) and determining viral titers by plaque assay on Vero-E6 cells. A549-ACE2 cells (ATCC) were plated at 1 × 10^6^ cells per well in a 6-well plate in DMEM (Corning) supplemented with HEPES (Corning), penicillin/streptomycin (Corning), GlutaMAX (Thermo Fisher Scientific), and 10% fetal bovine serum (FBS) (Sigma) the day before infection. The cells were then infected at MOI 0.1 for 1 h at 37°C, after which the overlying media was replaced with DMEM supplemented with HEPES, penicillin/streptomycin, GlutaMAX, and 2% FBS. After 48 h, cells were washed with PBS (Corning) and harvested using TrypLE (Life Technologies) before staining with Live/Dead Blue dead cell stain (Invitrogen) and a BV510-conjugated pan-HLA class I antibody (clone W6/32, BioLegend). Cells were fixed with 4% paraformaldehyde (Santa Cruz Biotechnology) and subsequently permeabilized with Perm/Wash Buffer (BD Biosciences). Intracellular staining was performed in Perm/Wash Buffer with an AF647-conjugated pan-HLA class I antibody (clone W6/32, BioLegend) and a SARS-CoV-2 nucleocapsid antibody (clone A20087H, mouse IgG2b isotype, BioLegend) followed by a secondary antibody stain with PE-Cy7-conjugated anti-mouse IgG2b (BioLegend). Fixed and stained cells were then washed and prepared for flow cytometry. All live virus experiments performed in the Ragon Institute Biosafety Level 3 (BSL3) laboratory following protocols approved by the Mass General Brigham Institutional Biosafety Committee. Flow cytometry was performed on a BD Symphony (BD Biosciences, San Jose, CA) and analyses were performed using FlowJo v10.7.1 and GraphPad Prism 9 software.

### Immunoprecipitation

HEK-293T cells were seeded at 2 X10^5^ in 6-well plates and, on the next day, were inoculated with ORF7a-expressing SCRPSY lentivirus stocks. At 40 hours post-infection, cells were detached by trypsinization and treated for 20 min with 10 mM methyl methanethiosulfonate (MMTS, Thermo Scientific) in PBS on ice. Cells were then lysed with ice-cold lysis buffer (25 mM Tris, pH 7.4, 150 mM NaCl, 0.4% NP-40 (Sigma), 5% glycerol, supplemented with 1X complete protease inhibitor (Roche). After lysis on ice for 20 min, followed by centrifugation at 10,000 rpm for 10 min at 4°C, clarified lysates were mixed with 2 μg mouse anti-ORF7a monoclonal antibody (Genetex), mouse anti-HLA-A (Proteintech 66013-1-Ig), or rabbit anti-HLA-B (ThermoFisher Scientific PA5-35345), or mouse anti-HLA-A, B, C (W6/32) and rotated with 30 μl pre-equilibrated Protein G Sepharose 4 Fast Flow resin (GE healthcare) for 2 hours at 4°C. The resin was then washed 3 times with PBS and the bound proteins were eluted with SDS-PAGE sample buffer and analyzed by Western blotting.

### Western blot analysis

Cell lysates and immunoprecipitates were separated on NuPage Novex 4-12% Bis-Tris Mini Gels (Invitrogen), and NuPAGE™ MES SDS running buffer (Invitrogen, NP0002) was used. Proteins were blotted onto nitrocellulose membranes. Thereafter, the blots were probed with primary antibodies and followed by secondary antibodies conjugated to IRDye 800CW or IRDye 680. Fluorescent signals were detected using an Odyssey scanner (LI-COR Biosciences).

To measure the sensitivity of MHC-I to Endo H treatment, A549 cells were transduced with lentivirus stocks expressing ORF7a proteins from SARS-CoV, SARS-CoV-2, or bat SARSr CoVs at an MOI of 0.5. After 40 hours, cells were treated with puromycin to remove untransduced cells and, after additional 20 hours, cells were harvested and treated with Endo H (New England Biolabs) for 60 minutes. The whole cell lysates were loaded on SDS-PAGE for Western blot analysis.

### HIV-1 Gag-specific T-cell clones

CD8+ T cell clones specific for the HLA-A11-restricted Gag epitope ACQGVGGPGHK (AK11) were isolated and expanded using a previously described protocol (33). Briefly, PBMCs from an HIV-infected individual were thawed, washed in warm X-VIVO-15, and resuspended at a concentration of 1 × 10^7^ cells/mL. PBMCs were stimulated for 3 hours with 10 μg/mL of Gag AK11 peptide and T cells producing IFN-γ in response were enriched using the IFN-γ Secretion Detection and Enrichment Kit (130-054-201; Miltenyi Biotec, Bergisch Gladbach, Germany) in accordance with the manufacturer’s instructions. Enriched T cells were plated at a series of dilutions in 96-well plates with irradiated feeder medium (RPMI 1640 supplemented with 10% FBS, l-glutamine, and PenStrep [R-10]) with 1 × 10^6^ cells/mL 5000 rad irradiated PBMC + 50 U/mL IL-2 + 10 ng/mL IL-15 (both from NCI BRB Preclinical Biologics Repository) + 0.1 μg/mL each of anti-CD3 (Ultra-LEAF purified anti-human CD3 antibody clone OKT3; BioLegend, San Diego, CA) and anti-CD28 (Ultra-LEAF purified Anti-human CD28 antibody clone 28.2; BioLegend), 37°C 5% CO2. One month later, wells were selected from the most dilute plate that showed growth (<1/3 of wells containing cell populations) and each was screened for specificity to Gag-AK11 peptide pool by CD107a staining and flow cytometry. Specific clones were expanded through biweekly stimulations with irradiated feeder medium. Clone specificity was further confirmed by peptide stimulation and CD107a staining on the day before performing T cell antigen recognition assays.

### T-cell antigen recognition assay

To measure antigen presentation, 293T cells were transduced with a retrovirus vector (LMNi-bsd) expressing HLA A11, or an empty vector control. The cells were subsequently transduced with empty SCRPSY or SCRPSY expressing an ORF7a protein at an MOI of 1. After selection in puromycin for 24 hours cells were transfected with an HIV-1 Gag expression plasmid. these ‘target’ cells were washed to remove the selection media, plated in 96 well plates and then cocultured with a Gag-AK11-specific CD8+ T-cell clone in RPMI 1640 medium supplemented with 10% fetal bovine serum (FBS), 2 mM l-glutamine, 100 units/ml penicillin, 100 μg/ml streptomycin, and 50 units/ml IL-2 (NCI BRB Preclinical Biologics Repository) at 37°C 5% CO2. After 16 hours, cells were pelleted and supernatants were harvested for IFN-γ ELISA. Cells were then washed in 2% FBS phosphate-buffered saline + 2mM EDTA and surface stained with fluorochrome-conjugated antibodies to CD3-Brilliant Violet 785 clone OKT3, CD8-BV605 clone SK1, MHC-I-PacBlue clone W6/32, (all from BioLegend), CD4-APC R700 clone RPA-T4, (BD), Gag KC57-FITC (Beckman Coulter) and Fixable Aqua Viability Dye (Invitrogen). Counting beads were also added. Samples were analyzed by flow cytometry, and data analysis was performed with FlowJo, version 10, software.

### IFN-γ ELISA

Levels of IFN-γ in supernatants from T cell clone recognition assays were measured by ELISA using the ELISA MAX Deluxe Set Human IFN-g kit (Biolegend) following the manufacturer’s instructions. Concentrations were determined by interpolating from a standard curve.

### Deconvolution Microscopy

Five thousand A549 cells were seeded onto gelatin (Millipore ES-0060B) coated 8-chamber #1.5 borosilicate glass bottom slides (LabTek 155409) and transduced with SCRPSY lentiviruses expressing ORF7a or a mutant thereof at an MOI of 1. For some wells 2 μl CellLight ER-GFP BacMam 2.0 (ThermoFisher C10590) was added at the time of infection. The cells were fixed at 48 h.p.i. with 4% formaldehyde (ThermoFisher 28908) in PBS (Thermo AM9624) for 15 min at RT. Cells were rinsed with PBS and blocked with 10% goat serum (Sigma G9023-10ML) in PBS for 20 min. Following 10 min permeabilization at room temperature with 0.2% IGEPAL CA-630 (Sigma I3021), 10% goat serum in PBS, cells were probed for 1 hr at RT with primary antibodies in 0.1% Tween20 (Fisher BP337-500), 10% goat serum in PBS as indicated: HLA-A (Sigma H1650-100STS, 0.4 μg/ml), beta-2 microglobulin (Novus NBP2-44471, 0.4 μg/ml) or ORF7a (Bioworld NCP011, 2 μg/ml). After three rinses with 0.1% Tween20 (Fisher BP337-500) in PBS, cells were probed with secondary antibodies for 45 min at RT in 0.1% Tween20, 10% goat serum, Hoechst 33258 (Abcam ab145596, 1 μg/mL) in PBS as indicated: goat anti-rabbit (Novus NBP1-72732C, 1.4 μg/ml), goat anti-mouse (Novus NBP1-72739JF646, 1.4 μg/ml or ThermoFisher Scientific A11029 4 μg/ml). Cells were rinsed twice with 0.1% Tween20 in PBS and twice with PBS and imaged by deconvolution microscopy (DeltaVision OMX SR imaging system). Image analysis was done using ImageJ (Version 2.0.0-rc-59/1.51w).

---

**Figure S1.**
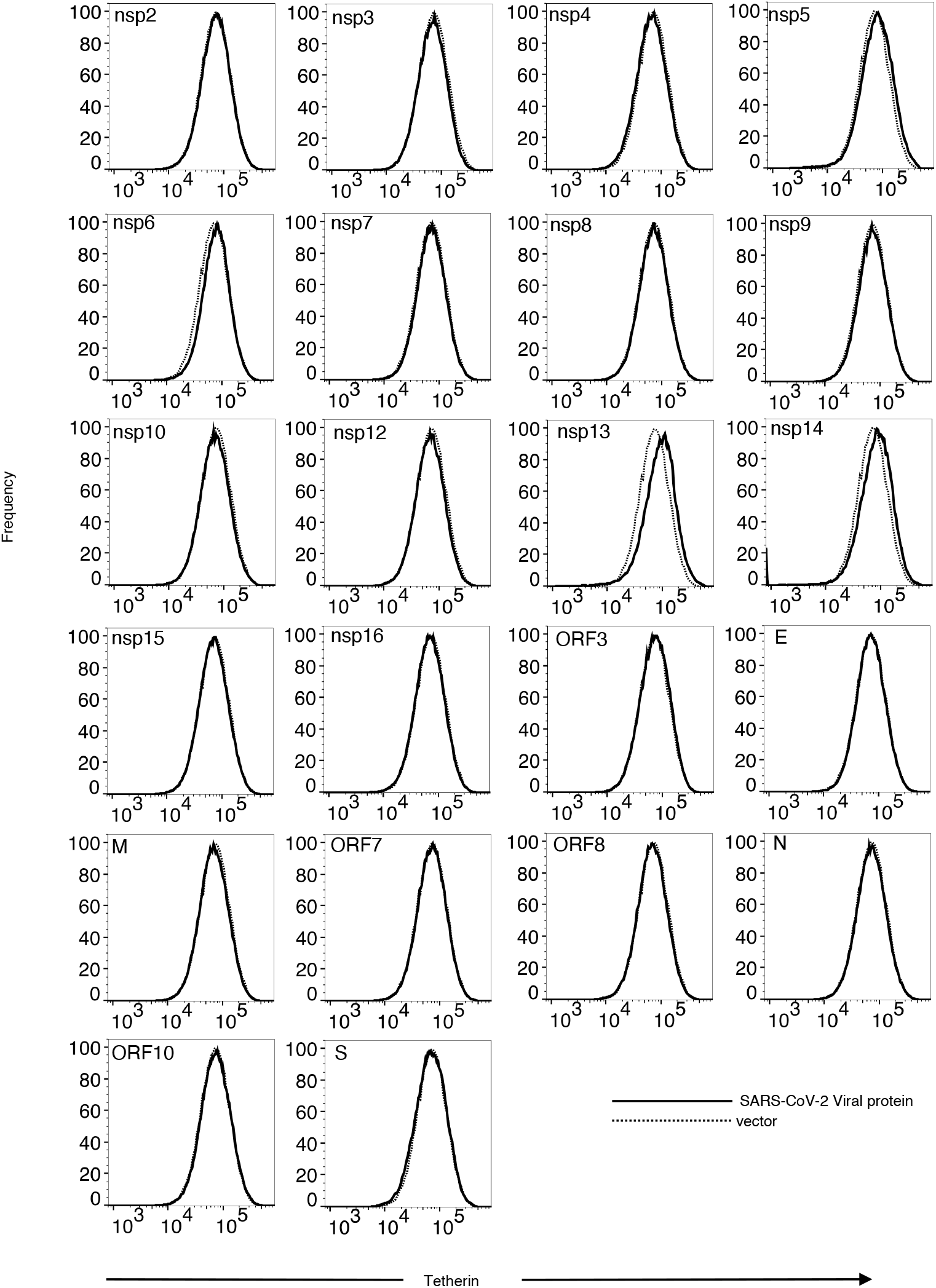
No effect of SARS-CoV-2 proteins on cell surface levels of Tetherin. Human 293T cells transduced with pSCRPSY-based lentiviral vector expressing individual SARS-CoV-2 viral proteins, at an MOI of 0.5 and cell surface Tetherin-HA measured with anti-HA followed and flow cytometry. Cells transduced (TagRFP+ population) with a viral protein expression vector (solid line) and empty lentiviral vector (dotted line) were gated and compared. Representative of two experiments.

**Figure S2.**
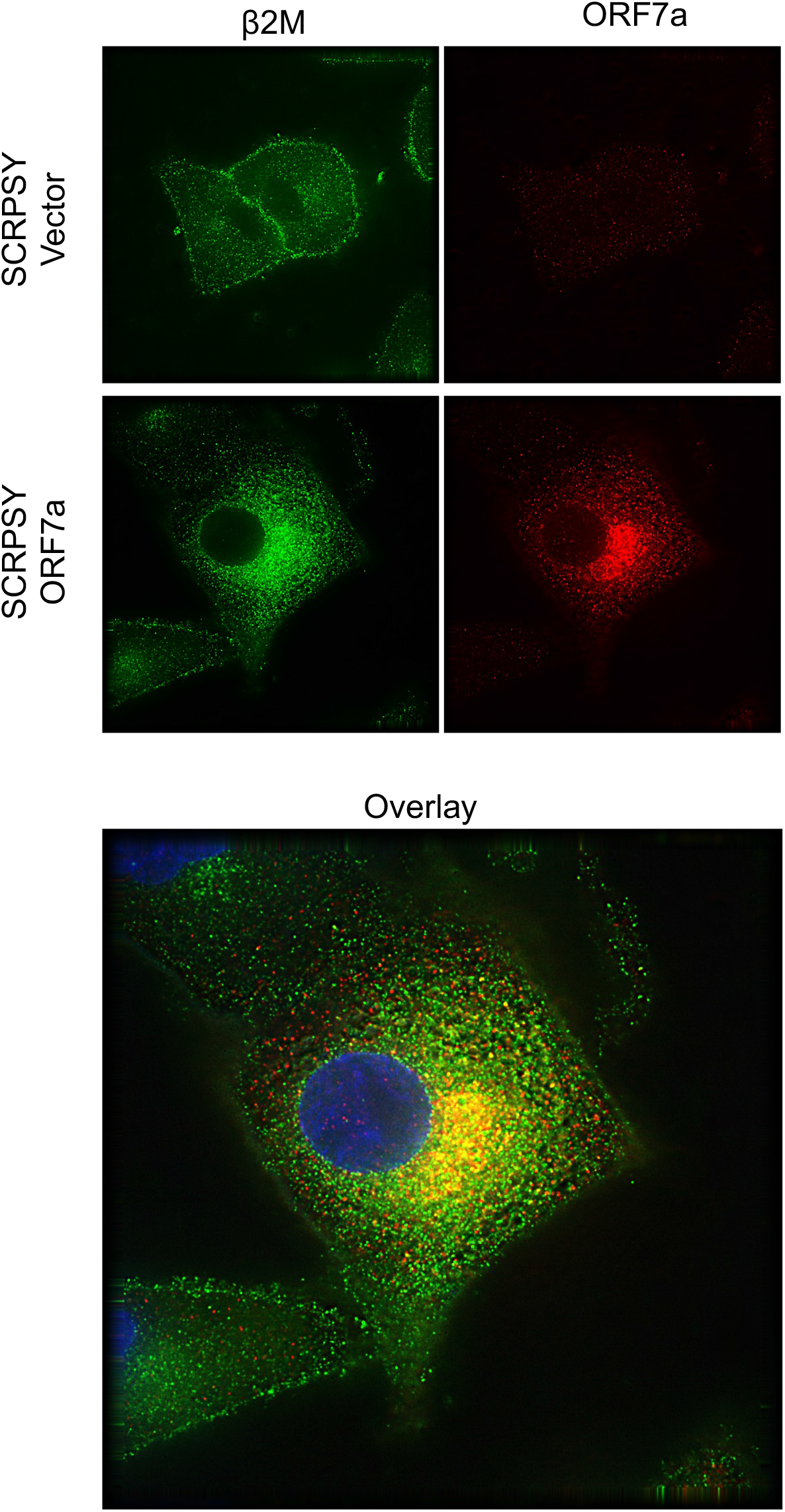
Redistribution of β2M in SARS-CoV-2 ORF7a-expressing cells. Immunofluorescent staining of A549 cells with anti-β2M (green) and anti-ORF7a following transduction with empty SCRPSY (vector) or ORF7a-expressing SCRPSY (ORF7a). A single optical section following deconvolution microscopy is shown.

**Figure S3.**
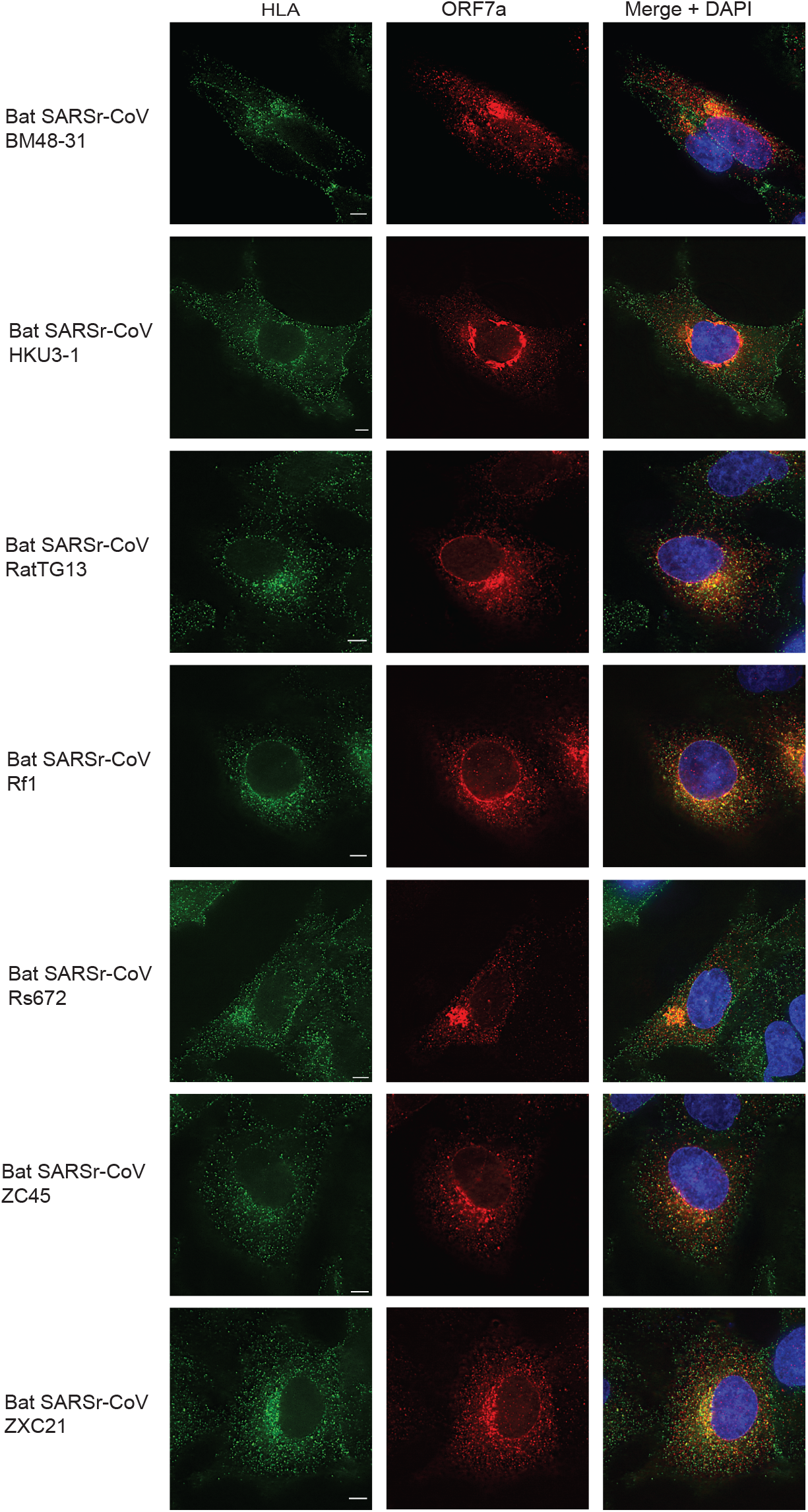
Distribution of MHC-I in bat SARSr-CoV ORF7a-expressing cells. Immunofluorescent staining of A549 cells with anti-HLA-A (green) and anti-ORF7a (red) following transduction with SCRPSY (vector) or SCRPSY expressing ORF7a from the indicated SARSr-CoV. A single optical section following deconvolution microscopy is shown.

**Figure S4.**
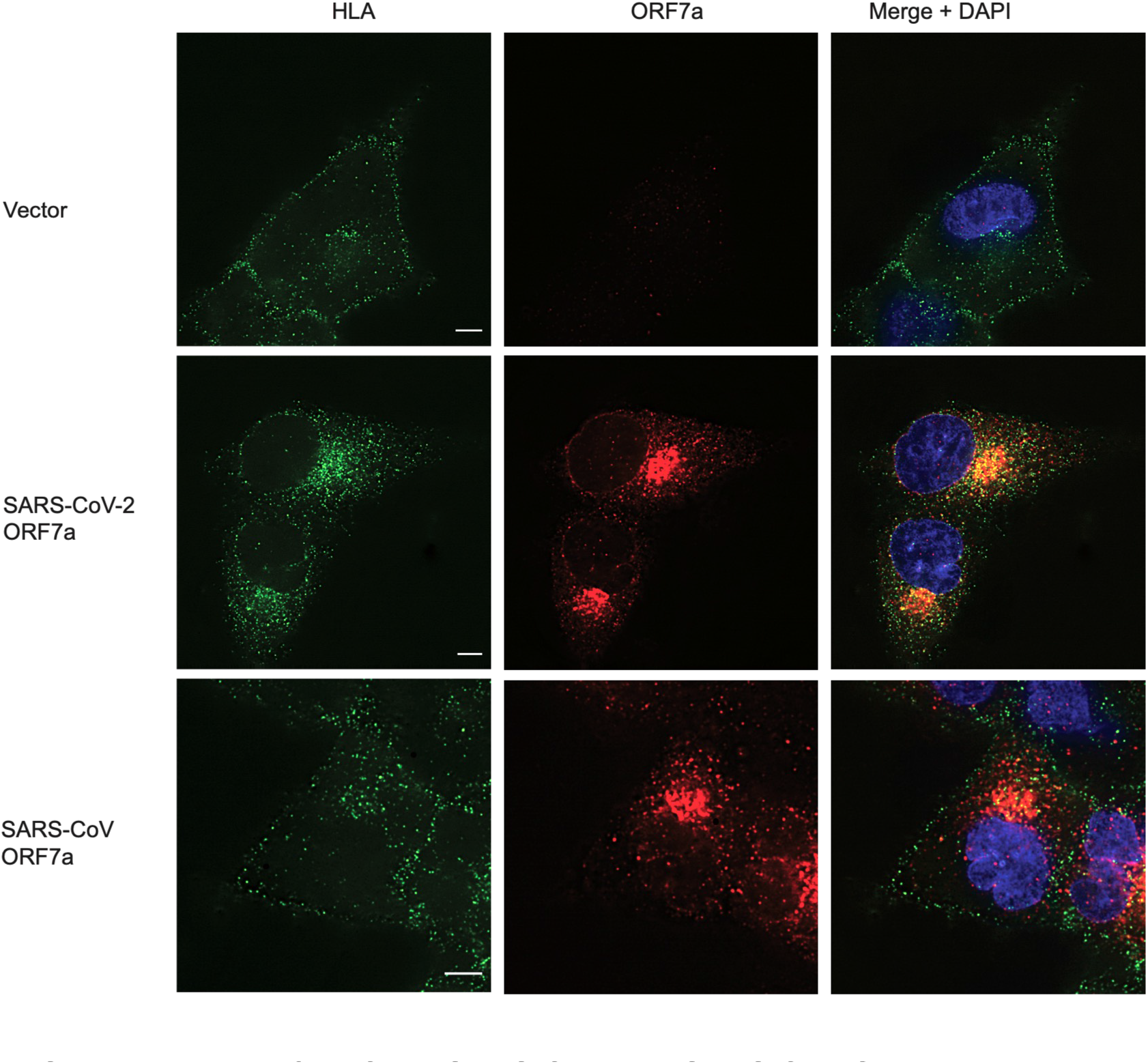
Distribution of MHC-I in SARS-CoV-2 or SARS-CoV ORF7a-expressing cells. Immunofluorescent staining of A549 cells with anti-HLA-A (green) and anti-ORF7a (red) following transduction with SCRPSY (vector) or ORF7a-expressing SCRPSY. A single optical section following deconvolution microscopy is shown.

## Notes

### Competing Interest Statement

The authors have declared no competing interest.

