## Supplementary Information Table for "Inhibition of major histocompatibility complex-I antigen presentation by sarbecovirus ORF7a proteins"

Supplementary Information Table: Antibodies and reagents used in this paper

Immunoblotting (IB)

| Target Antigen | Target species | Host species | Clone | Company | Catalog # |
| --- | --- | --- | --- | --- | --- |
| SARS-CoV / SARS-CoV-2 ORF7a |  | Mouse | 3C9 | GeneTex | GTX632602 |
| ORF7a |  | Rabbit | polyclonal | Bioworld | NCP0011 |
| HLA-A | Human | Rabbit | EP1395Y | abcam | ab52922 |
| HLA-A, B, C | Human | Mouse | EMR8-5 | abcam | ab70328 |
| Calreticulin | Human | Rabbit | polyclonal | abcam | ab2907 |
| ERp57 | Human | Rabbit | polyclonal | abcam | ab10287 |
| TAP1 | Human | Rabbit | polyclonal | ThermoFisher Proteintech | 11114-1-AP |
| TAP2 | Human | Rabbit | polyclonal | abcam | ab180611 |
| TAP2 | Human | Rabbit | polyclonal | ThermoFisher Invitrogen | PA5-37414 |
| Tapasin | Human | Rabbit | polyclonal | ThermoFisher Invitrogen | PA5-42731 |
| Tapasin | Human | Mouse | TO-3 | Santa Cruz | sc-80647 |
| Tubulin | Human | Mouse | DM1A | Sigma | T9026 |

Flow cytometry (FCM)

| Target Antigen | Target species | Host species | Clone | Conjugation | Company | Catalog # |
| --- | --- | --- | --- | --- | --- | --- |
| HLA-A, B, C | Human | Mouse | W6/32 | AF488 | Biolegend | 311413 |
| HLA-A, B, C | Human | Mouse | W6/32 | AF647 | Biolegend | 311414 |
| HLA-A2 | Human | Mouse | BB7.2 | APC | Biolegend | 343308 |
| HLA-A | Human | Rat | YTH 862.2 |  | abcam | Ab00200-8.1 |
| HA epitope | tag | Mouse | 16B12 | AF488 | Biolegend | 901509 |
| HA epitope | tag | Mouse | 16B12 | APC | Biolegend | 901524 |

Immunoprecipitation (IP)

| Target Antigen | Target species | Host species | Clone | Company | Catalog # |
| --- | --- | --- | --- | --- | --- |
| SARS-CoV / SARS-CoV-2 ORF7a |  | Mouse | 3C9 | GeneTex | GTX632602 |
| HLA-A, B, C | Human | Mouse | W6/32 | Biolegend | 311402 |
| HLA-A, B, C | Human | Mouse | 5C5B7 | Proteintech | 66013-1-Ig |
| HLA-B | Human | Rabbit | polyclonal | ThermoFisher Invitrogen | PA5-35345 |

Immunofluorescence (IF)

| Target Antigen | Target species | Host species | Clone | Company | Catalog # |
| --- | --- | --- | --- | --- | --- |
| SARS-CoV / SARS-CoV-2 ORF7a |  | Rabbit |  | Bioworld | NCP0011 |
| HLA-A, B, C | Human | Mouse | W6/32 | Sigma | H1650-100TST |
| HLA-A | Human | Rabbit | EP1395Y | Abcam | ab52922 |
| B2M | Human | Mouse | B2M/961 | Novus | NBP2-44471 |
| IgG H+L | Rabbit | Goat |  | Novus | NBP1-72732C |
| IgG H+L | Mouse | Goat |  | Novus | NBP1-72739JF646 |
| IgG H+L | Mouse | Goat |  | Thermo | A11029 |
